# Protein-protein interaction network topology as a measure of bacterial plasmid-chromosome co-evolution

**DOI:** 10.1101/610774

**Authors:** Nghia T.H. Tran, Arun Decano, Tim Downing, Alexander D. Rahm

## Abstract

In this work, we investigate the evolvability of plasmids by examining the topology of plasmid-chromosome gene interactions in Escherichia coli ST131. We discover a convergence of the ratio of non-trivial loops per protein-protein interaction, which allows us to introduce a new invariant of bacterial PPINs: the indirect connectivity value.

## 1 Introduction

Antimicrobial resistance (AMR) is a major public health concern driving infection by extra-intestinal pathogenic bacteria, including Escherichia coli ST131-O25b [1]. ST131 has extensive non-susceptibility to a wide range of antibiotic classes as a result of its acquistion of key AMR genes encoded on plasmids ([2], [3]), permitting it to adapt to treatments [4].

Plasmids are circular DNA molecules capable of self-replication and typically of transmission by conjugation between cells ([5], [6], [7]). They can be classified based on their incompatibility (Inc) group, and for ST131 most commonly they are in IncF [8]. Although plasmids in ST131 typically encode genes for post-segregation killing and stable inheritance to ensure their propagation, they can be lost or may recombine with other plasmids in the same cell ([9], [10], [11]). As a result of this mixing and their extensive array of mobile genetic elements (MGEs), plasmids may rearrange extensively even within a clonal radiation [12].

Plasmids may increase ST131 fitness through the gain of AMR genes, but equally may impair cell reproduction due to the energetic cost of plasmid replication and maintenance. In an environment with antibiotics driving AMR but with opportunities for plasmid conjugation and recombination, there is potential for plasmid-chromosome co-evolution to optimise gene dosage, coordinate expression and recalibrate virulence and pathogenicity. Protein-protein interactions (PPIs) are a major component of this adaptive process and so numbers of plasmid-chromosome PPIs are likely to be positively correlated with time [13]. Bacterial PPI networks (PPINs) are functionally rich sources of information on protein relationships ([14], [15], [16]). PPIN connectivity thus acts as a proxy for co-evolvability: more connected protein pairs co-evolve more so than distantly connected ones.

Signatures of chromosome-plasmid co-evolution in PPINs can be more informative if they incorporate both direct and indirect connectivity across proteins. Topological data analysis (TDA) has been used to investigate complex and high-dimensional biological datasets, like breast cancer genomes[17]. In this study, we use TDA to quantify the higher dimensional network structure in E. coli ST131-related plasmids by quantifying rates of “holes” (in technical terms, non-trivial loops) in PPINs. From the ratio of “holes” per interaction, we derive a new indicator, the *indirect connectivity value*, which is positively correlated with higher connectivity. Using chromosomal PPINs as a comparator, we find that specific plasmids’ proteins interact more with chromosomal proteins, and observe that the ratio of “holes” per interaction converges across interaction thresholds to our indirect connectivity value; hence the latter can be used as a stable unbiased metric. This highlights potential for it as a measure of plasmid-chromosome co-evolution that is independent of PPIN size, and could be applied to predict the capability of unobserved plasmid-chromosome combinations to consider future potential AMR repertoires in pathogens.

## 2 Methods

### 2.1. Plasmid data sources

Here, we examine plasmids common in E. coli genomes: pCA14, pEK204, pEK499, pEK516, pEC958, pJIE186-2 and pSE15 (Table 1). Particularly prevalent among ST131 are pEK204, pEK499 and pEK516, and have been informative when examining plasmid diversity in ST131 (Lanza et al 2014). All had genes allowing stable plasmid inheritance and post-segregation killing. IncFII plasmid pEK499 lacks a traX gene for conjugation and is associated the historical acquisition of a blaCTX-M-15 gene in ST131 that is tightly correlated with its pandemic nature [18]. Another blaCTX-M-15-positive plasmid (pEK516) considered here is structurally similar to pEK499 with about 75 percent similarity. Similar to these is pEC958 from a UK blaCTX-M-15-positive UTI that has 85 percent identity with pEK499 and is missing the latter’s second tra region (transfer genes for conjugation) due to an IS26-mediate blaTEM-1 insertion (Phan et al 2015, Forde et al 2014). We scrutinise pEK204 too because it has a tra region adequate for conjugation, though some tra genes may be lost during culturing (Woodford et al). As a contrast in terms of AMR, we use pJIE186-2 that has numerous virulence factor genes that are chromosomally encoded in some E. coli, but has no known AMR genes and incomplete genes for conjugation (Zong et al 2013). We also compare these to conjugative plasmids pCA14 (blaCTX-M-15-positive, Li et al 2015) and pECSF1 (aka pSE15), which is from a commensal isolate to serve as a genetically distinct outgroup (O150:H5, fimH41) (Toh et al 2010).

**Table 1.** The seven plasmids’ properties and relevant genetic characteristics for ST131 AMR and potential chromosomal PPIs. The IncF1A replicon in pJIE186-2 is partial.

| Plasmid name | Accession ID | Total genes | Length (kb) | Inc group | Conjugative (tra region) | Antibiotic resistance |
| --- | --- | --- | --- | --- | --- | --- |
| pCA14 | CP009231 | 181 | 155.4 | F2/F1A/F1B | Yes | Yes |
| pEK204 | EU935740 | 112 | 93.7 | I1 | Yes | Yes |
| pEK499 | EU935739 | 185 | 117.5 | F2/F1A | No | Yes |
| pEK516 | EU935738 | 103 | 64.6 | F2A | No | Yes |
| pEC958A | HG941719 | 142 | 135.6 | F1A/F2 | No | Yes |
| pJIE186-2 | NC_020271 | 138 | 137.7 | F1B/F2A/F1A | No | No |
| pECSF1 | NC_013655 | 150 | 122.3 | F2A/FIB | Yes | No |

### 2.2. Bacterial PPIN construction

We extracted E. coli K12 MG1655 PPIN data from the Search Tool for the Retrieval of Interacting Genes/Proteins (STRING) database [19] v10 (Szklarczyk et al 2019) via the R v3.5.2 packages BiocManager v1.30.4, dplyr v0.8.0.1, genbankr v1.10.0, rentrez v1.2.1, STRINGdb v1.22.0, tidyverse v1.2.1, and VennDiagram v1.6.20. Using a combined score threshold of at least 400 to remove spurious interactions, this dataset had 4,146 distinct protein-coding genes, of which 4,121 had interaction information, resulting in 105,457 interactions. The STRING database combined scores were used as the primary metric to determine valid interactions, because they integrate multiple types of evidence while controlling for random interactions [19].

For each dataset or plasmid, we obtained the number of pairwise PPIs within that unit and with the chromosome for unique genes. We used the above mentioned R packages to examine the connectivity of the plasmid and chromosomal gene products across the STRING combined score values from 400 to 900. Proteins with informative interactions were identical in pEK499 and pEK516, so only the former was examined further (Table 1).

We used results of a previous PPIN clustering analysis (Table 1 of Miryala et al [20]) as a control to test our approach. This had five clusters with *n* = 60 genes in total, which were examined as a unit, as was the cluster 1 subset of *n* = 30 proteins and the pan-cluster *n* = 19 AMR-related proteins from all five clusters to allow inter-and intra-cluster patterns to be examined (Table 2). This showed that the within-group interactions per protein (3.8–4.6) and chromosomal interactions per protein (8.2–14.9) were consistent across groups and were subsequently comparable to values for the chromosome and plasmids.

**Table 2.** The numbers of protein-coding genes per Miryala et al [20] gene set. All were unique and had interaction data (bar six in the first set of 60). The numbers of interactions within each cluster is shown, followed by the numbers per group with the chromosome.

| Subset name | Total genes | Number of interactions within-group | Number of interactions with chromosome | Number of interactions per protein - plasmid | Number of interactions per protein - chromosome |
| --- | --- | --- | --- | --- | --- |
| All | 60 | 229 | 2,307 | 3.8 | 10.1 |
| Group 1 | 30 | 138 | 1,125 | 4.6 | 8.2 |
| All AMR | 19 | 57 | 852 | 4.0 | 14.9 |

### 2.3. PPIN topology determined by the Vietoris–Rips complex

The Vietoris–Rips complex was introduced independently by Vietoris and Rips [21]. We constructed the Vietoris–Rips complex on our data given a threshold *N* for the minimum PPI combined score. The set of proteins are the vertex set, connected by edges if an interaction is present. For each triple (*A, B, C*) of proteins, if the interaction score of the three contained pairs (*A, B*), (*B, C*) and (*C, A*) are all at least the threshold, then we fill in the empty (1-dimensional, 1D) triangle of edges which we already have drawn to connect these three pairs, by a solid (2D) triangle with corners *A, B* and *C*. For each quadruple (*A, B, C, D*) of proteins, if the interaction values of the six contained pairs (*A, B*), (*B, C*), (*C, D*), (*A, C*), (*B, D*) and (*D, A*) are all at least the threshold, then we fill in the empty (2D) tetrahedral surface which we already have drawn as the solid triangles (*A, B, C*), (*B, C, D*), (*C, D, A*) and (*D, A, B*), with a solid (3D) tetrahedron of corners *A, B, C* and *D*. We continue in higher dimension *m* on each (*m* + 1)-tuple (*A*_1_, *A*_2_, …, *A*_*m*+1_), if the interaction scores of the contained pairs are all at least the threshold, then we fill in the already created (*m* − 1)-dimensional hypersurface with an *m*-dimensional standard simplex.

There are publicly available implementations of the Vietoris–Rips complex, for instance in the SAGE computer algebra system. However, due to the high number of interactions in the chromosomal dataset explored here (105,457), general purpose implementations were not efficient enough, so we used specialised software to take into account the properties of our type of datasets, and to optimise the Betti number computations needed to be computed on the Vietoris–Rips complex. This software developed by one of the authors (Rahm) [22] uses the sparsity of the boundary matrices to process the rank computations efficiently in LinBox [23], and a hard-coded dimensional truncation of the Vietoris–Rips complex to avoid the high number of higher-dimension simplices that would be obtained for the full chromosomal dataset. For each dataset, we computed the three first Betti numbers of the Vietoris–Rips complex from STRING combined scores of 400 to 900 with a step size of 25.

## 3 Results

We examined 4, 146 chromosomal gene products’ PPIN topology across 105, 457 pairwise PPIs in E. coli using conventional methods as a contrast for indirect connectivity information derived from TDA. With the above software detailed in the Methods, we can overcome the complexity of the protein interactions graph to compute the first Betti number of the Vietoris–Rips complex, which counts *non-trivial* loops. A loop is *non-trivial* if it cannot be completely filled in with solid triangles from the Vietoris–Rips complex (see Methods for their construction). As a consequence, a non-trivial loop is composed of at least four proteins whose interactions constitute a closed chain such that not every pair of proteins has an interaction. Thus, a non-trivial loop represents a “hole” in the direct interactions, indicating indirect interations between proteins. For each considered interaction score threshold *N*, we compute the ratio

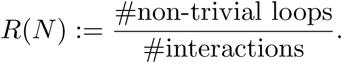

We present our topological results in Table 3 for the plasmid pEK204, in Table 4 for the plasmid pEK499, and in Figures 1 to 2b for the ratio *R*(*N*) plotted over the threshold *N*. The number of full chromosomal PPIs declines nearly linearly from 105, 457 to 14, 555 when the score threshold becomes more stringent (from 400 to 900), namely with a correlation coefficient square of *r*^2^ = 0.964 against the linear variation of the threshold, and similarly the ratio of non-trivial loops per interaction drops nearly linearly from *R*(400) = 0.518 to *R*(900) = 0.35, namely with *r*^2^ = 0.905 against the linear variation of the threshold.

**Table 3.** PPIN TDA information for plasmid pEK204 in relation to the E. coli chromosome across STRING combined score threshold that range from strict (900) to relaxed (400). For pEK204, the number of loops and loops/interaction declined considerably with rising score, indicating less connectivity.

| Threshold | Triangles | Interactions | Proteins | Two dimensional bubbles | Non-trivial loops | Connected components | Non-trivial loops per interaction |
| --- | --- | --- | --- | --- | --- | --- | --- |
| 900 | 8 | 58 | 57 | 4 | 1 | 4 | 0.017241379 |
| 875 | 8 | 73 | 72 | 4 | 2 | 5 | 0.02739726 |
| 850 | 10 | 84 | 81 | 5 | 3 | 5 | 0.035714286 |
| 825 | 10 | 102 | 98 | 5 | 6 | 7 | 0.058823529 |
| 800 | 10 | 112 | 109 | 5 | 6 | 8 | 0.053571429 |
| 775 | 12 | 126 | 122 | 6 | 7 | 9 | 0.055555556 |
| 750 | 16 | 148 | 138 | 8 | 9 | 7 | 0.060810811 |
| 725 | 17 | 171 | 159 | 8 | 12 | 9 | 0.070175439 |
| 700 | 17 | 196 | 183 | 8 | 13 | 9 | 0.066326531 |
| 675 | 17 | 215 | 202 | 8 | 14 | 10 | 0.065116279 |
| 650 | 17 | 233 | 219 | 8 | 15 | 10 | 0.064377682 |
| 625 | 19 | 251 | 236 | 9 | 15 | 10 | 0.059760956 |
| 600 | 19 | 266 | 250 | 9 | 16 | 10 | 0.060150376 |
| 575 | 19 | 302 | 283 | 9 | 19 | 10 | 0.062913907 |
| 550 | 19 | 323 | 301 | 9 | 21 | 9 | 0.06501548 |
| 525 | 19 | 344 | 320 | 9 | 22 | 8 | 0.063953488 |
| 500 | 19 | 379 | 349 | 9 | 27 | 7 | 0.071240106 |
| 475 | 23 | 406 | 371 | 11 | 30 | 7 | 0.073891626 |
| 450 | 24 | 459 | 412 | 11 | 38 | 4 | 0.082788671 |
| 425 | 24 | 497 | 447 | 11 | 41 | 4 | 0.08249497 |
| 400 | 27 | 561 | 496 | 13 | 54 | 3 | 0.096256684 |

**Table 4.** PPIN TDA information for plasmid pEK499 in relation to the E. coli chromosome across STRING combined score threshold that range from strict (900) to relaxed (400). For pEK499, the number of loops and loops/interaction was stable irrespective of the score, indicating consistent connectivity, unlike pEK204.

| Threshold | Triangles | Interactions | Proteins | Two dimensional bubbles | Non-trivial loops | Connected components | Non-trivial loops per interaction |
| --- | --- | --- | --- | --- | --- | --- | --- |
| 900 | 2 | 68 | 68 | 1 | 11 | 12 | 0.16176471 |
| 875 | 2 | 88 | 84 | 1 | 16 | 13 | 0.18181818 |
| 850 | 2 | 97 | 93 | 1 | 16 | 13 | 0.16494845 |
| 825 | 5 | 113 | 106 | 1 | 16 | 13 | 0.14159292 |
| 800 | 9 | 119 | 112 | 3 | 15 | 14 | 0.12605042 |
| 775 | 9 | 135 | 129 | 3 | 15 | 15 | 0.11111111 |
| 750 | 9 | 146 | 138 | 3 | 15 | 13 | 0.10273973 |
| 725 | 8 | 180 | 168 | 3 | 20 | 13 | 0.11111111 |
| 700 | 8 | 215 | 193 | 3 | 31 | 14 | 0.14418605 |
| 675 | 12 | 238 | 211 | 5 | 33 | 13 | 0.13865546 |
| 650 | 12 | 277 | 245 | 5 | 36 | 11 | 0.1299639 |
| 625 | 16 | 311 | 272 | 7 | 37 | 7 | 0.11897106 |
| 600 | 16 | 332 | 291 | 7 | 37 | 5 | 0.11144578 |
| 575 | 22 | 375 | 326 | 10 | 41 | 4 | 0.10933333 |
| 550 | 26 | 407 | 353 | 12 | 44 | 4 | 0.10810811 |
| 525 | 34 | 445 | 379 | 16 | 52 | 4 | 0.11685393 |
| 500 | 36 | 486 | 417 | 17 | 54 | 4 | 0.11111111 |
| 475 | 44 | 530 | 453 | 21 | 57 | 3 | 0.10754717 |
| 450 | 51 | 586 | 495 | 24 | 67 | 3 | 0.11433447 |
| 425 | 55 | 646 | 545 | 26 | 74 | 2 | 0.11455108 |
| 400 | 66 | 758 | 636 | 31 | 89 | 2 | 0.11741425 |

**Figure 1.**
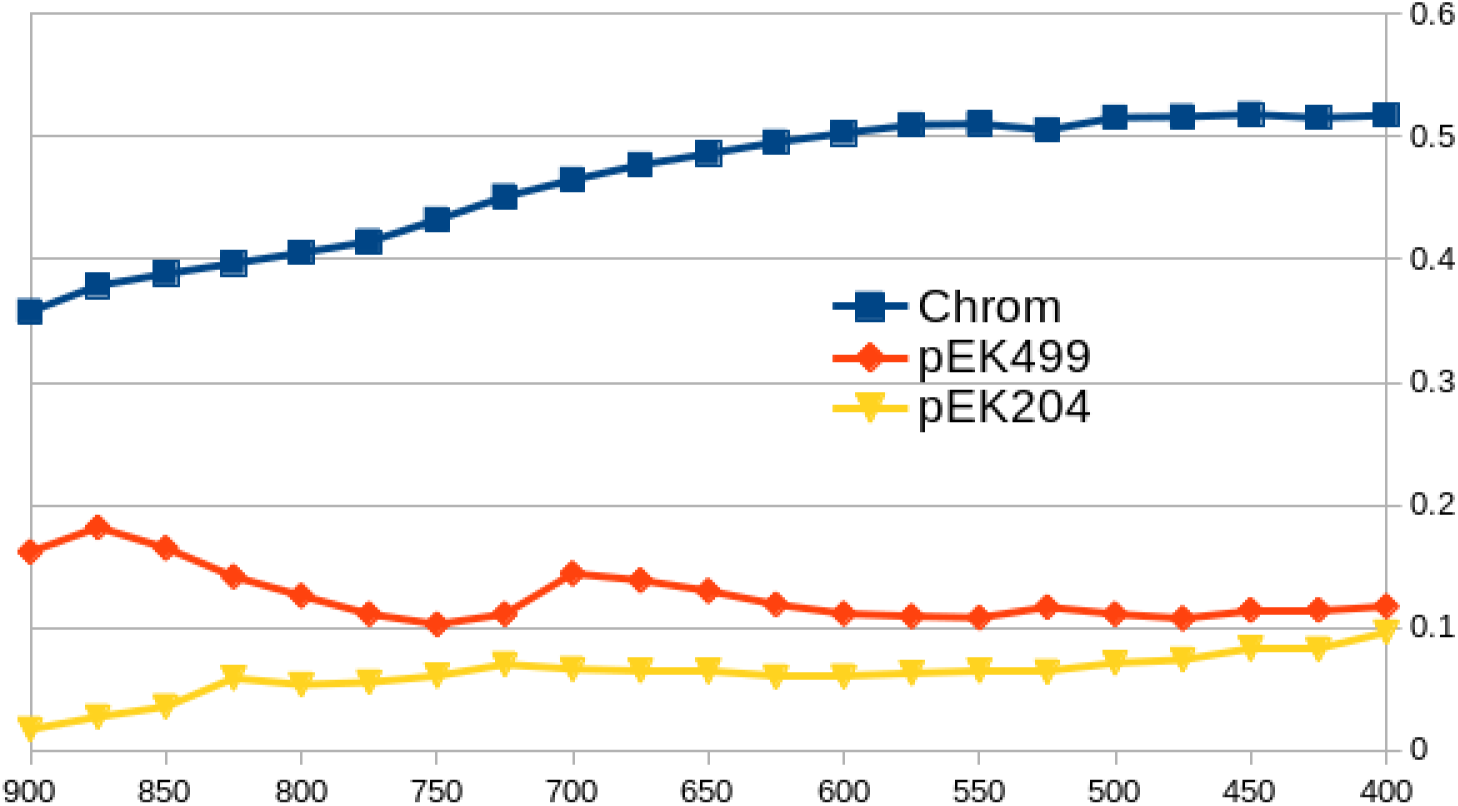
Full E. coli ST131 genome, Plasmid pEK204, Plasmid pEK499: (Non-trivial loops)/interaction plotted over threshold

**Figure 2.**
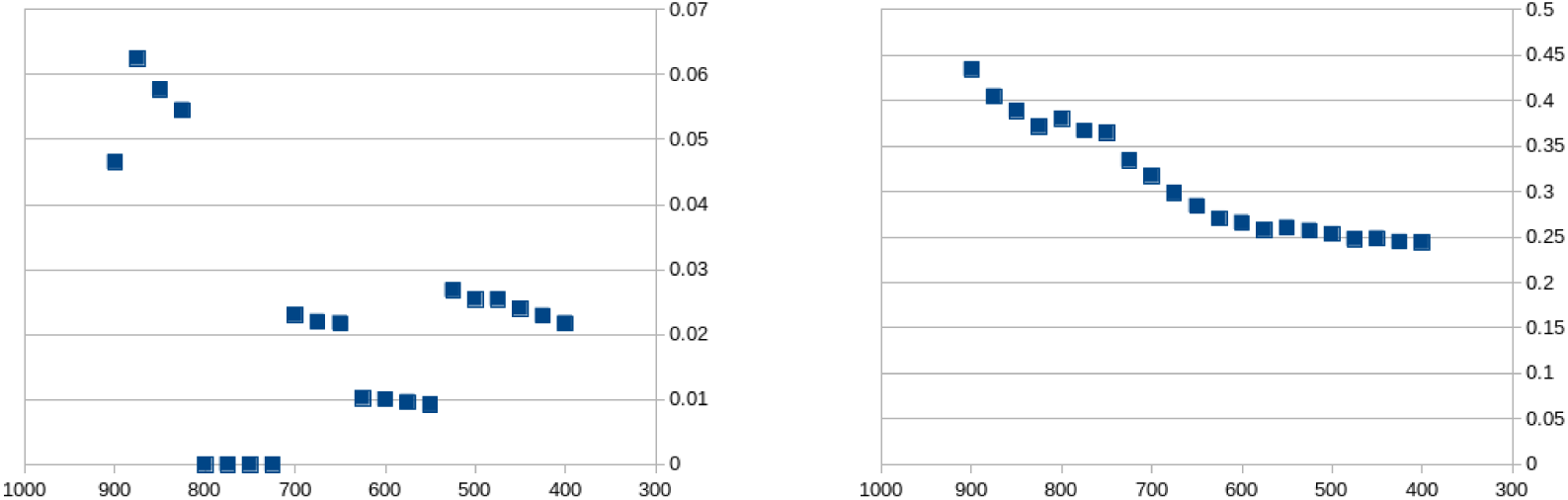
(Non-trivial loops)/interaction plotted over threshold. **(a)** Data for 30 genes from Miryala et al Table 1, Group 1 only **(b)** Data for 60 genes from Miryala et al Table 1, all Groups 1-5

## 4 Discussion

The main observation of this paper is that on the chromosomal proteins of E. coli ST131 bacteria and on their common plasmid proteins pEK204, pEK499 and pEK516, the ratio *R*(*N*) of non-trivial loops per interaction *converges to some value close to its value at the threshold N* = 400, when we are varying the threshold *N* down towards 400. This phenomenon is displayed in Figure 1.

### 4.1. A new measure of indirect connectivity in PPINs

Based on this observation, whenever we have convergence, we interpret *R*(400) as a new *indirect connectivity value* (ICV). As we are interested in the plasmid datasets, we use the full chromosome dataset with 4,146 proteins as a baseline. On the latter, we measure *R*(400) = 0.518, and we record this as our baseline value *b*_*fc*_. Then for a plasmid set, we define the *indirect connectivity* to be the ratio

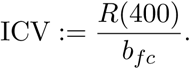

The reason for our choice of name is the following. For a loop to be non-trivial, at least four genes need to be involved, and while their interactions constitute a closed chain, not between every pair of them can we have an interaction. In other words, there is a “hole” of missing direct interactions, facing the presence of indirect interactions along the chain. The higher the indirect connectivity value is, the better the chance to connect two genes which are not directly interacting, by a chain of interactions passing via other genes. We collect its values in Table 5.

**Table 5.** Indirect connectivity values, compared with the ratio 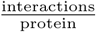. Note that the latter ratio is approximately monotonic when varying from threshold 900 down to threshold 400.

| Dataset | ICV | $\frac{\text{interactions}}{\text{protein}}$ at threshold 900 | $\frac{\text{interactions}}{\text{protein}}$ at threshold 400 |
| --- | --- | --- | --- |
| genomes 4,146 | 100% | 4.48 | 25.2 |
| pEK204 | 18.6% | 1.02 | 1.13 |
| pEK499 | 22.7% | 1.00 | 1.19 |
| pJIE186-2 | 0% | 0.86 | 0.97 |

### 4.2. Observations on the individual plasmids

The number of loops and loops/interaction connecting the pEK499 proteins to the chromosomal ones is stable irrespective of the STRING combined score, indicating consistent connectivity (Table 4). In contrast, the same metrics for pEK204 show a continual decline, showing less connectivity as the score threshold increases (Table 3). Although all plasmids’ gene proteins were substantially less connected to the chromosomal proteins, the pattern for pEK499 was distinct. This was emphasised by the difference across the other plasmids in Table 6, showing that the proteins of pCA14, pEC958A and pECSF1 had no chromosomal interactions, and that pJIE186-2 had relatively few. This indicates that pEK204, pEK499 and pEK516 may share a stronger co-evolutionary signal with the E. coli chromosome, and that this is not dependent on conjugative ability, Inc group, nor plasmid size as per Table 1.

**Table 6.**
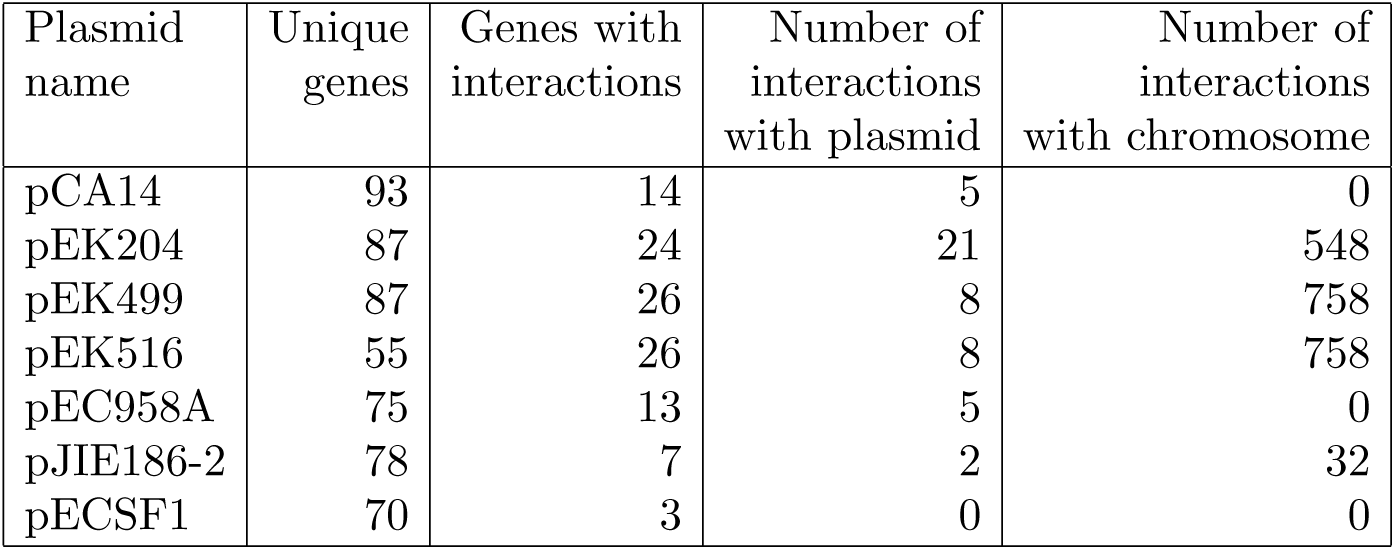
The numbers of unique protein-coding genes per plasmid that had interactions with other proteins given a combined score threshold of 400, followed by the number of interactions within the plasmid and with the chromosome.

| Plasmid name | Unique genes | Genes with interactions | Number of interactions with plasmid | Number of interactions with chromosome |
| --- | --- | --- | --- | --- |
| pCA14 | 93 | 14 | 5 | 0 |
| pEK204 | 87 | 24 | 21 | 548 |
| pEK499 | 87 | 26 | 8 | 758 |
| pEK516 | 55 | 26 | 8 | 758 |
| pEC958A | 75 | 13 | 5 | 0 |
| pJIE186-2 | 78 | 7 | 2 | 32 |
| pECSF1 | 70 | 3 | 0 | 0 |

### 4.3. Future work

We hypothesise that the adaptive benefit associated with antibiotic resistance genes encoded on pEK204, pEK499 and pEK516 may be the driver of long-term co-existance and co-evolution of these plasmids with the E. coli ST131 chromsome [4]. To explore this further, plasmid-chromosome interactions in other Enterobacteriaceae species are needed to investigate if the ICV (Table 5) reflects PPIN hierarchy and co-evolutionary patterns more broadly.

## 5 Conclusion

We evaluate the structure of chromosome-plasmid PPINs in bacterial genomes to detect signatures of indirect connectivity potentially associated with co-evolution between the vertically inherited chromosomes and plasmids that can be transferring horizontally. Plasmid conjugation, recombination and AMR gene repertoire are key aspects in the spread of pandemic E. coli ST131. Our indirect connectivity value (ICV) is derived from TDA to detect indirect connectivity and so provides a new classificiation method applicable to other bacterial species and plasmids. By showing that the proteins on plasmids pEK204, pEK499 and pEK516 are more connected to the E. coli chromosome, we highlight new avenues for investigating AMR.

